# Prevalence, characteristics, and clinical significance of concomitant cardiomyopathies in subjects with bicuspid aortic valve

**DOI:** 10.1101/496919

**Authors:** Hyeonju Jeong, Chi Young Shim, Darae Kim, Jah Yeon Choi, Kang-Un Choi, Soo Youn Lee, Geu-Ru Hong, Jong-Won Ha

## Abstract

**Background:** In this study, the prevalence, characteristics, and clinical significance of concomitant specific cardiomyopathies were investigated in subjects with bicuspid aortic valve (BAV).

**Methods:** We retrospectively evaluated 1,186 adults with BAV (850 males, mean age 56 ± 14 years). Left ventricular non-compaction, hypertrophic cardiomyopathy, and idiopathic dilated cardiomyopathy were diagnosed when patients fulfilled current clinical and echocardiographic criteria. Clinical and echocardiographic characteristics including comorbidities, heart failure presentation, BAV morphology, function, and aorta phenotypes in BAV subjects with or without specific cardiomyopathies were compared.

**Results:** Overall, 67 subjects (5.6 %) had concomitant cardiomyopathies: 40 (3.4%) patients with left ventricular non-compaction, 17 (1.4%) with hypertrophic cardiomyopathy, and 10 (0.8%) with dilated cardiomyopathy. BAV subjects with hypertrophic cardiomyopathy had a higher prevalence of diabetes mellitus, heart failure with preserved ejection fraction, and tended to have type 0 phenotype, while BAV subjects with dilated cardiomyopathy showed a higher prevalence of chronic kidney disease and heart failure with reduced ejection fraction. BAV subjects with left ventricular non-compaction were significantly younger, predominantly male, and had greater BAV dysfunction and a higher prevalence of normal aorta shape. In multiple regression analysis, presence of cardiomyopathy was independently associated with heart failure (odds ratio 2.866, 95% confidential interval 1.652–4.974, p < 0.001) even after controlling confounding factors.

**Conclusion:** Concomitant cardiomyopathies were observed in 5.6% of subjects with BAV. A few clinical and echocardiographic characteristics including comorbidities, heart failure presentation, BAV morphology, function, and presence of aortopathy were found. The presence of cardiomyopathy was independently associated with heart failure.

## Introduction

Bicuspid aortic valve (BAV) is the most common congenital heart valve disease, with an overall incidence of approximately 1% in the general population [1, 2]. Subjects with BAV often present with aortic dilatation and may have mechanical functional alterations in vasculatures [3, 4]. In addition, BAV is a highly heritable trait, often associated with other congenital heart defects or genetic syndromes [5, 6].

Regarding myocardial characteristics in subjects with BAV, most previous studies have focused on subclinical left ventricular dysfunction associated with increased aortic stiffness [7-9]. Although a possible association between BAV and specific cardiomyopathies (CMs) based on common genetic traits has been proposed in several case reports [10, 11], data on the prevalence of coexisting specific CMs in subjects with BAV are limited. In a previous study, an 11% incidence of left ventricular non-compaction (LVNC) in 109 patients with BAV was reported [12]. In a recent large population study, an only 0.4% prevalence of hypertrophic CM (HCM) was reported in 5,430 patients with BAV, similar to the general population [13]. However, the prevalence of concomitant specific CMs might be different based on ethnicity. Moreover, data are lacking regarding the clinical and echocardiographic characteristics based on the type of concomitant CMs. Therefore, in the present study, the prevalence, characteristics, and clinical significance of concomitant CM including LVNC, HCM, and idiopathic dilated CM (DCM) were determined using a large Korean BAV registry.

## Methods

### Study Population

We retrospectively reviewed the subjects diagnosed with BAV using transthoracic echocardiography in Severance Cardiovascular Hospital from January 2003 to December 2017. A total of 1,186 subjects (850 males, mean age 56 ± 14 years) were included in this study. All echocardiographic studies in subjects with BAV were manually reviewed for confirmation. Patient medical data as recorded by the physicians were carefully reviewed by two experienced observers who were blinded to echocardiography results. The institutional review board of Severance Hospital approved the present study, which was conducted in compliance with the Declaration of Helsinki. The subjects were classified into four groups based on the presence of specific CM.

Standard two-dimensional and Doppler measurements were performed following American Society of Echocardiography guidelines [14]. BAV was diagnosed based on anatomic evaluation of the aortic valve, when only two cusps were unequivocally identified in systole and diastole in the short-axis view and with a clear “fish mouth” appearance during systole [15]. The BAV morphology was classified into four types based on position and pattern of raphe and cusps. Type 1 indicated fusion of the left coronary and right coronary cusps, type 2 indicated the fusion of the right coronary and noncoronary cusps, and type 3 indicated the fusion of the left coronary and noncoronary cusps. Type 0 was defined when there were two developed cusps and no raphe (true type) [15, 16]. The severity of aortic stenosis or aortic regurgitation was assessed using integrated approaches [17, 18]. The dimensions of the sinus of Valsalva, sinotubular junction, and ascending aorta were measured as previously described [7, 15]. Presence of aortopathy was defined as ascending aorta dimension ≥ 40 mm, as previously defined.

HCM was clinically diagnosed based on the presence of unexplained myocardial hypertrophy (wall thickness ≥ 15 mm) in the absence of local or systemic etiologies capable of producing the extent of hypertrophy evident [13]. Mild systemic hypertension was not an exclusion criterion in the diagnosis of HCM. Subjects with coexisting HCM were then subdivided into three morphologically obstructive HCM subgroups, non-obstructive HCM, non-apical HCM, or non-obstructive apical HCM, based on either the presence of left ventricular outflow track obstruction or the predominant hypertrophy at the left ventricular apex. LVNC was diagnosed based on the previously suggested echocardiographic criteria [19-21] and/or ratios between noncompacted and compacted layers of the left ventricular wall on cardiac magnetic resonance imaging [22]. If the diagnosis of LVNC was suspected but not confirmed using echocardiography, cardiac magnetic resonance imaging was performed based on clinician decision. DCM was defined as an ejection fraction < 40% in the presence of increased left ventricular dimension. Subjects with ischemic heart disease, uncorrected or corrected significant aortic stenosis or aortic regurgitation (≥ moderate degree), and other reversible causes were excluded from diagnosis of idiopathic DCM in this study. Echocardiographic data were gathered and analyzed by experienced sonographers blinded to each patient’s clinical data. Heart failure was diagnosed using current diagnostic criteria [23] and was categorized based on left ventricular ejection fraction (LVEF) as follows: heart failure with preserved ejection fraction (LVEF ≥ 50%) and heart failure with reduced ejection fraction (LVEF < 50%).

Continuous variables were expressed as mean ± standard deviation (SD). Categorical variables were expressed as number (percentage). Comparisons between the groups were performed using standard λ^2^ tests for categorical variables and paired *t*-tests for continuous variables. Univariate and multivariate logistic regression analyses were performed. All statistical analyses were performed using SPSS Statistics, software version 23.0 (IBM, Armonk, NY, USA); a p-value < 0.05 was considered statistically significant.

## Results

### Prevalence of coexistent CMs in BAV subjects

Overall, 67 subjects (5.6%) had concomitant CMs; 10 (0.8%) subjects with DCM, 17 (1.4%) with HCM, and 40 (3.4%) with LVNC. Among the subjects with coexistent HCM, five had obstructive HCM, six presented with non-obstructive HCM, and six with non-obstructive apical HCM (Fig 1).

**Fig 1.**
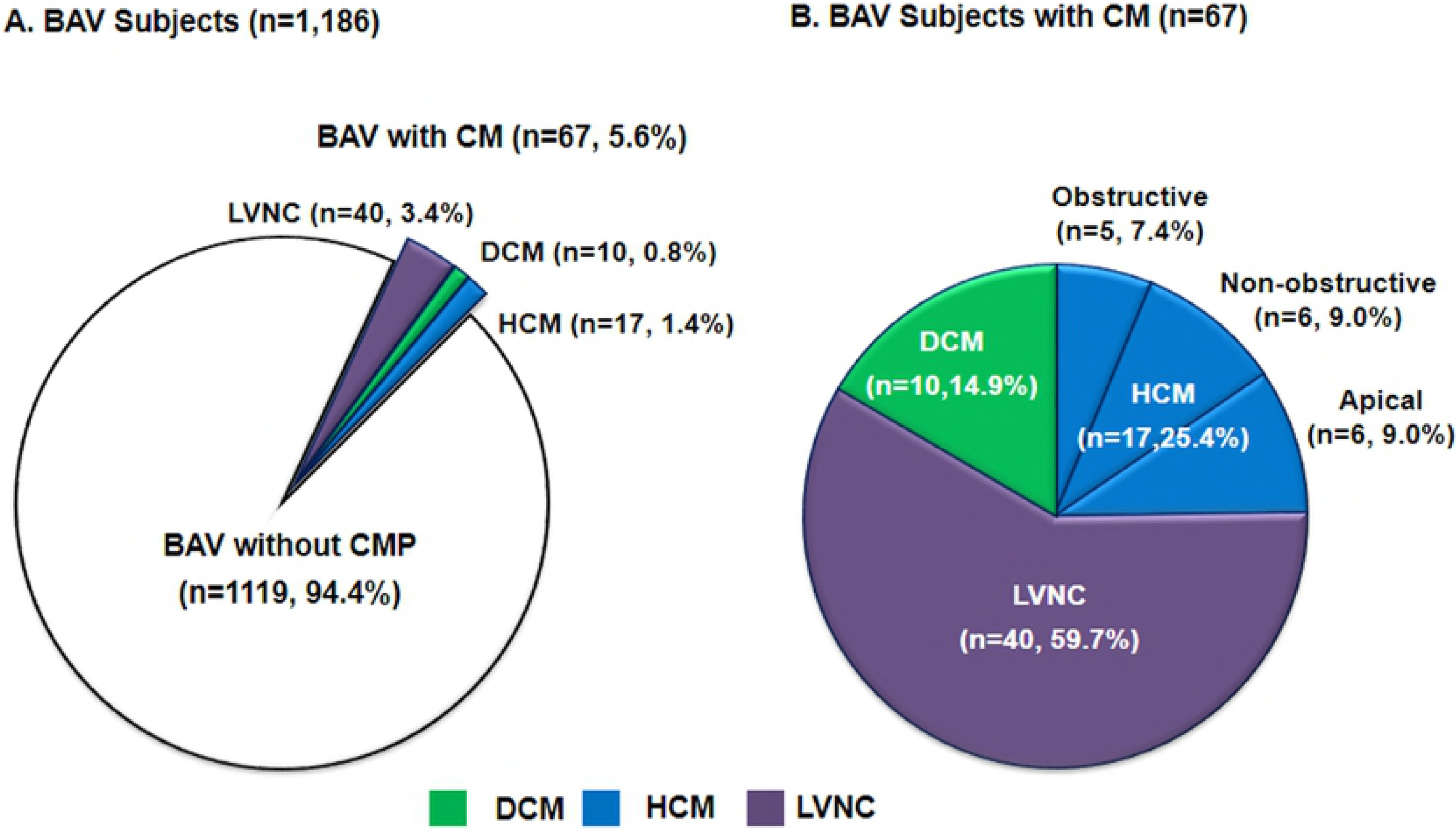
Prevalence of specific cardiomyopathies in BAV subjects

### Characteristics of BAV subjects with HCM, LVNC, or DCM

Baseline characteristics of the subjects with or without specific CMs are shown in Table 1. Subjects with DCM had a higher prevalence of chronic kidney disease and heart failure with reduced ejection fraction compared with those without CM. Subjects with HCM showed a higher prevalence of diabetes mellitus and heart failure with preserved ejection fraction than those without CM. Subjects with LVNC were younger and predominantly male compared with those without CM. In addition, subjects with LVNC exhibited a lower prevalence of heart failure than patients with DCM or HCM.

**Table 1.**
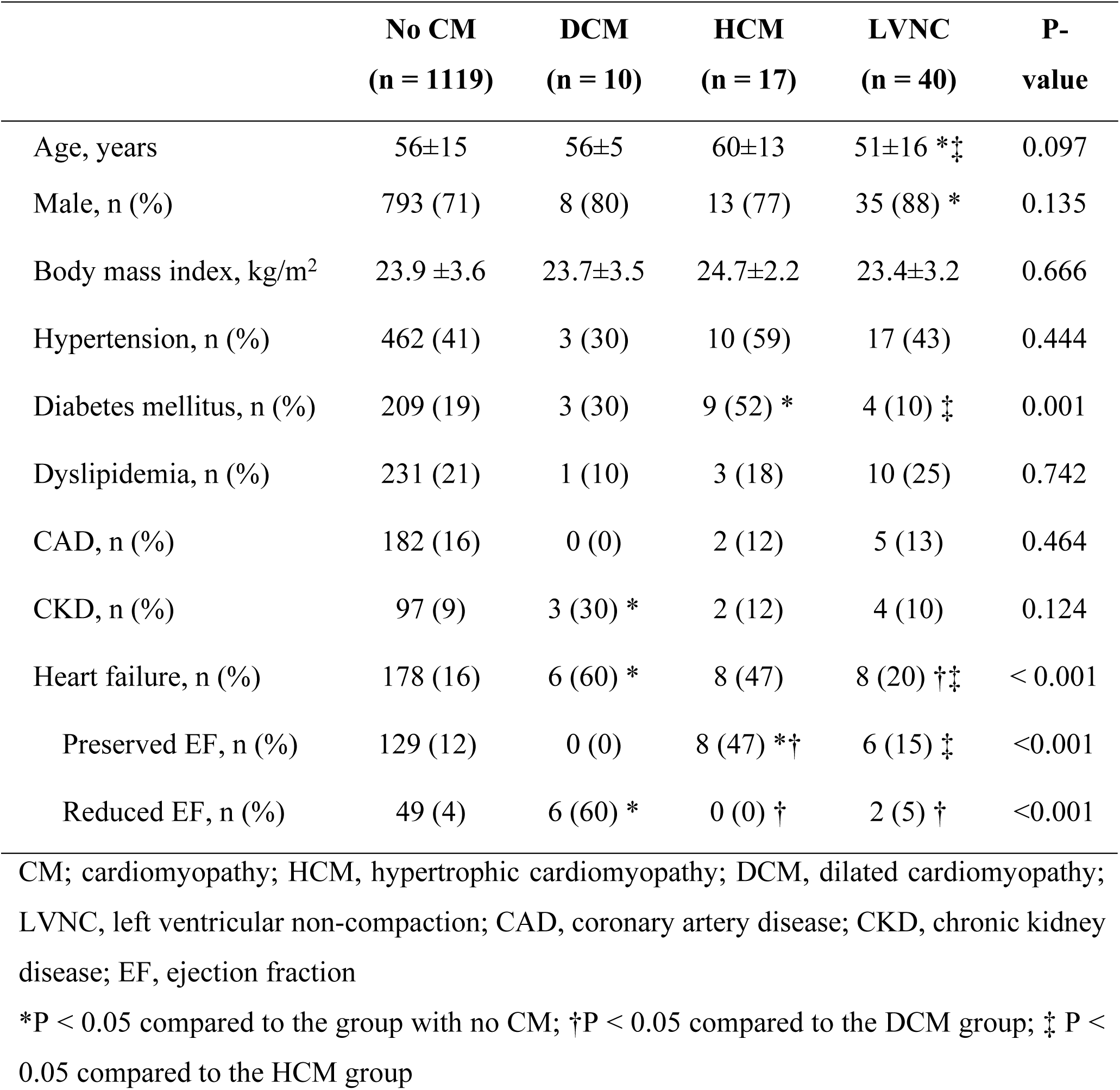
Baseline characteristics according to the presence of specific cardiomyopathy

Table 2 shows the structural and functional characteristics of the left ventricle in each group. The echocardiographic variables including left ventricle dimension, wall thickness, and LVEF were significantly different among the groups because of their own disease characteristics. LA volume index in BAV subgroups with specific CMs was significantly greater than in the BAV group without CMs. Early diastolic mitral annular tissue (e’) velocity and ratio of early diastolic mitral inflow velocity to e’ velocity (E/e’) in BAV subgroups with HCM and DCM were also significantly greater than in BAV subgroups without CMs. However, right ventricular systolic pressure was not significantly different among the groups.

**Table 2.** Left ventricle characteristics according to the presence of specific cardiomyopathy

|  | No CM<br>(n = 1119) | DCM<br>(n = 10) | HCM<br>(n = 17) | LVNC<br>(n = 40) | P-value |
| --- | --- | --- | --- | --- | --- |
| LVEDD, mm | 53.5±9.1 | 68.7±10.3* | 50.7±9.9† | 59.0±10.2*†‡ | <0.001 |
| LVESD, mm | 36.3±8.7 | 59.2±9.8* | 31.9±8.1*† | 41.9±9.5*†‡ | <0.001 |
| IVS thickness, mm | 10.7±2.2 | 10.1±2.6 | 14.7±4.2*† | 10.7±2.3‡ | <0.001 |
| PW thickness, mm | 10.5±2.0 | 10.3±2.2 | 12.2±2.3*† | 10.7±2.0‡ | 0.005 |
| LV ejection fraction, % | 63±12 | 29±11* | 70±8*† | 58±12*†‡ | <0.001 |
| LA volume index, ml/m<sup>2</sup> | 32.0±16.4 | 46.9±21.4* | 36.0±14.4 | 37.7±16.4* | 0.004 |
| e' velocity, cm/s | 6.1±2.5 | 3.5±1.4* | 4.1±1.6* | 6.7±3.0†‡ | <0.001 |
| E/e' | 13.0±5.9 | 20.0±6.6* | 17.5±7.3* | 12.8±5.4†‡ | <0.001 |
| RVSP, mmHg | 29±11 | 33±12 | 25±5 | 29±10 | 0.526 |

CM; cardiomyopathy; HCM, hypertrophic cardiomyopathy; DCM, dilated cardiomyopathy; LVNC, left ventricular non-compaction; LVEDD, left ventricular end-diastolic dimension; LVESD, left ventricular end-systolic dimension; IVS, interventricular septum; PW, posterior wall; LV, left ventricle; LA, left atrium; e', early diastolic mitral annular tissue; E/e', ratio of early diastolic mitral inflow velocity to early diastolic mitral annular tissue velocity; RVSP, right ventricular systolic pressure
\*P < 0.05 compared to the group with no CM; †P < 0.05 compared to the DCM group; ‡ P < 0.05 compared to the HCM group

The BAV characteristics in Table 3 and Figure 2 show that the type 1 BAV phenotype (fusion of right and left coronary cusps) was the most prevalent morphology in all groups. Although subjects with HCM tended to have type 0 phenotype, there was no statistically significant differences in BAV phenotypes among the groups. Subjects with DCM showed a higher prevalence of no or mild dysfunction, while subjects with LVNC exhibited a lower prevalence of no or mild dysfunction. Regarding the aorta phenotypes, subjects with LVNC revealed a higher prevalence of normal shape than those without CMs. Although statistical significance was not observed, more DCM patients had predominant AA phenotype, and none showed predominant sinus of Valsalva. Figure 3 illustrates the representative cases of coexisting CMs in subjects with BAV.

**Table 3.**
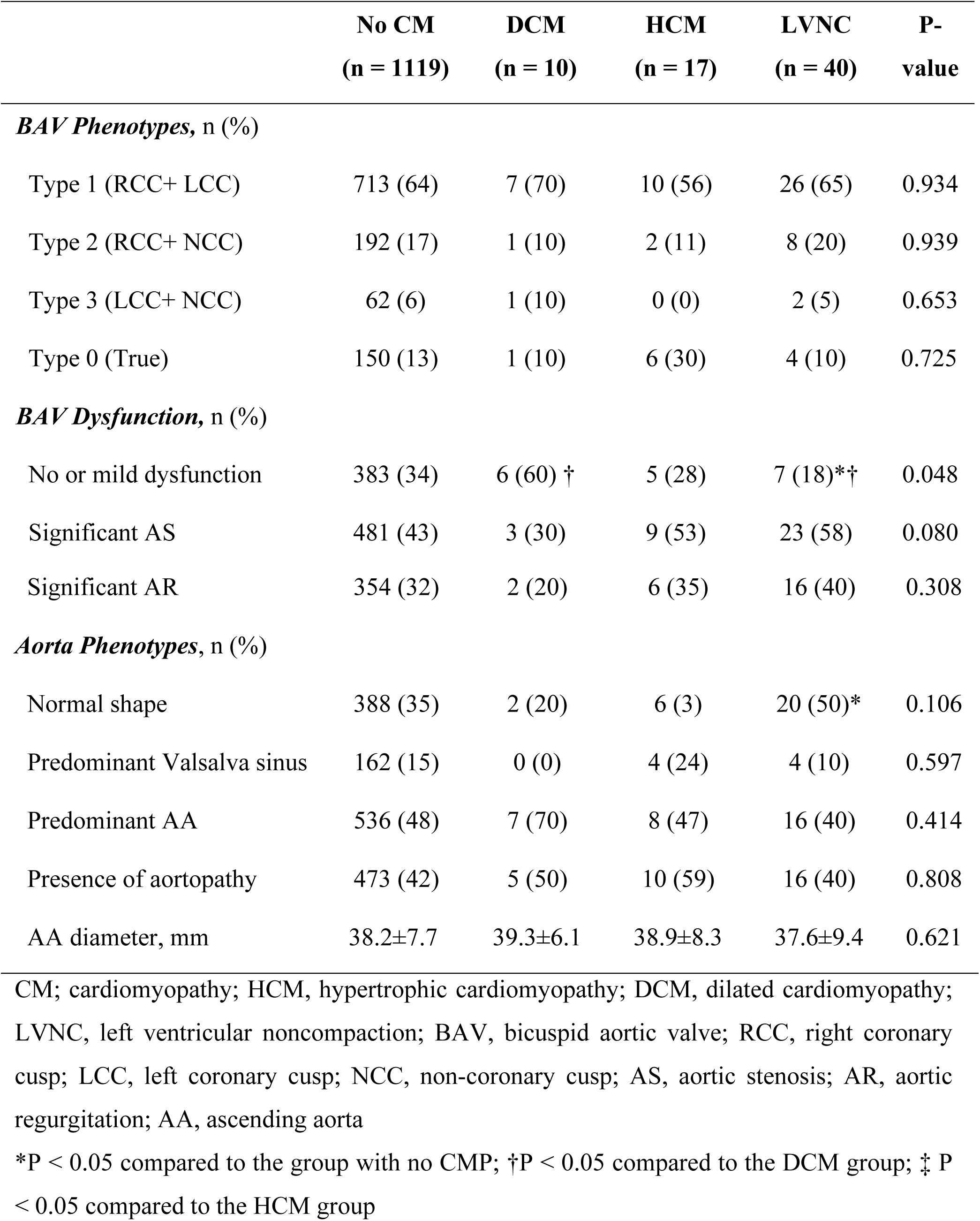
BAV and aorta characteristics according to the presence of specific cardiomyopathy

|  | **No CM**<br>**(n = 1119)** | **DCM**<br>**(n = 10)** | **HCM**<br>**(n = 17)** | **LVNC**<br>**(n = 40)** | **P-**<br>**value** |
| --- | --- | --- | --- | --- | --- |
| ***BAV Phenotypes, n (%)*** |  |  |  |  |  |
| Type 1 (RCC+ LCC) | 713 (64) | 7 (70) | 10 (56) | 26 (65) | 0.934 |
| Type 2 (RCC+ NCC) | 192 (17) | 1 (10) | 2 (11) | 8 (20) | 0.939 |
| Type 3 (LCC+ NCC) | 62 (6) | 1 (10) | 0 (0) | 2 (5) | 0.653 |
| Type 0 (True) | 150 (13) | 1 (10) | 6 (30) | 4 (10) | 0.725 |
| ***BAV Dysfunction, n (%)*** |  |  |  |  |  |
| No or mild dysfunction | 383 (34) | 6 (60) † | 5 (28) | 7 (18)*† | 0.048 |
| Significant AS | 481 (43) | 3 (30) | 9 (53) | 23 (58) | 0.080 |
| Significant AR | 354 (32) | 2 (20) | 6 (35) | 16 (40) | 0.308 |
| *Aorta Phenotypes, n (%)* |  |  |  |  |  |
| Normal shape | 388 (35) | 2 (20) | 6 (3) | 20 (50)* | 0.106 |
| Predominant Valsalva sinus | 162 (15) | 0 (0) | 4 (24) | 4 (10) | 0.597 |
| Predominant AA | 536 (48) | 7 (70) | 8 (47) | 16 (40) | 0.414 |
| Presence of aortopathy | 473 (42) | 5 (50) | 10 (59) | 16 (40) | 0.808 |
| AA diameter, mm | 38.2±7.7 | 39.3±6.1 | 38.9±8.3 | 37.6±9.4 | 0.621 |

CM; cardiomyopathy; HCM, hypertrophic cardiomyopathy; DCM, dilated cardiomyopathy; LVNC, left ventricular noncompaction; BAV, bicuspid aortic valve; RCC, right coronary cusp; LCC, left coronary cusp; NCC, non-coronary cusp; AS, aortic stenosis; AR, aortic regurgitation; AA, ascending aorta
\*P < 0.05 compared to the group with no CMP; †P < 0.05 compared to the DCM group; ‡ P < 0.05 compared to the HCM group

**Fig 2.**
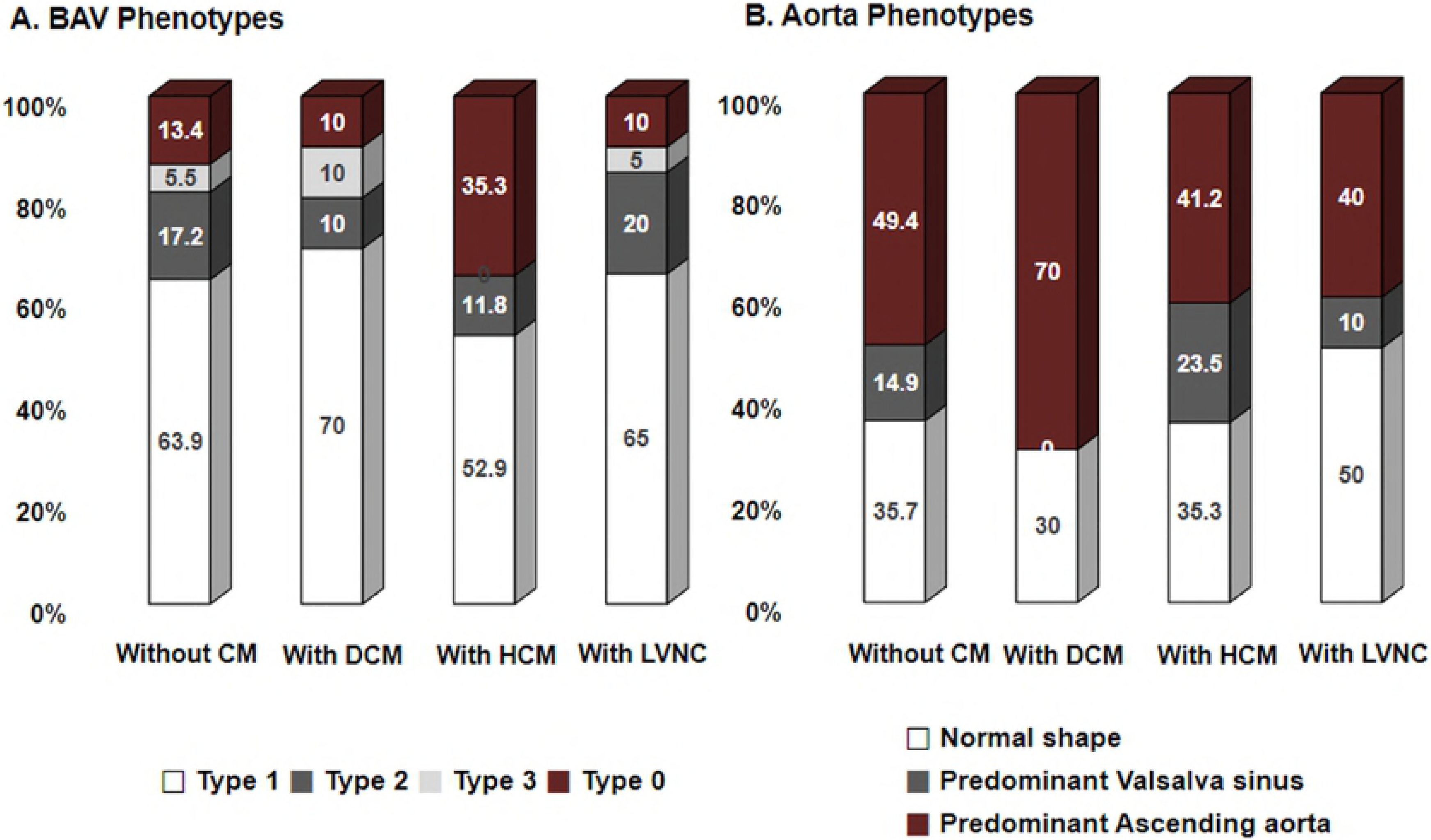
BAV phenotypes and aorta phenotypes according to the presence of specific cardiomyopathies

**Fig 3.**
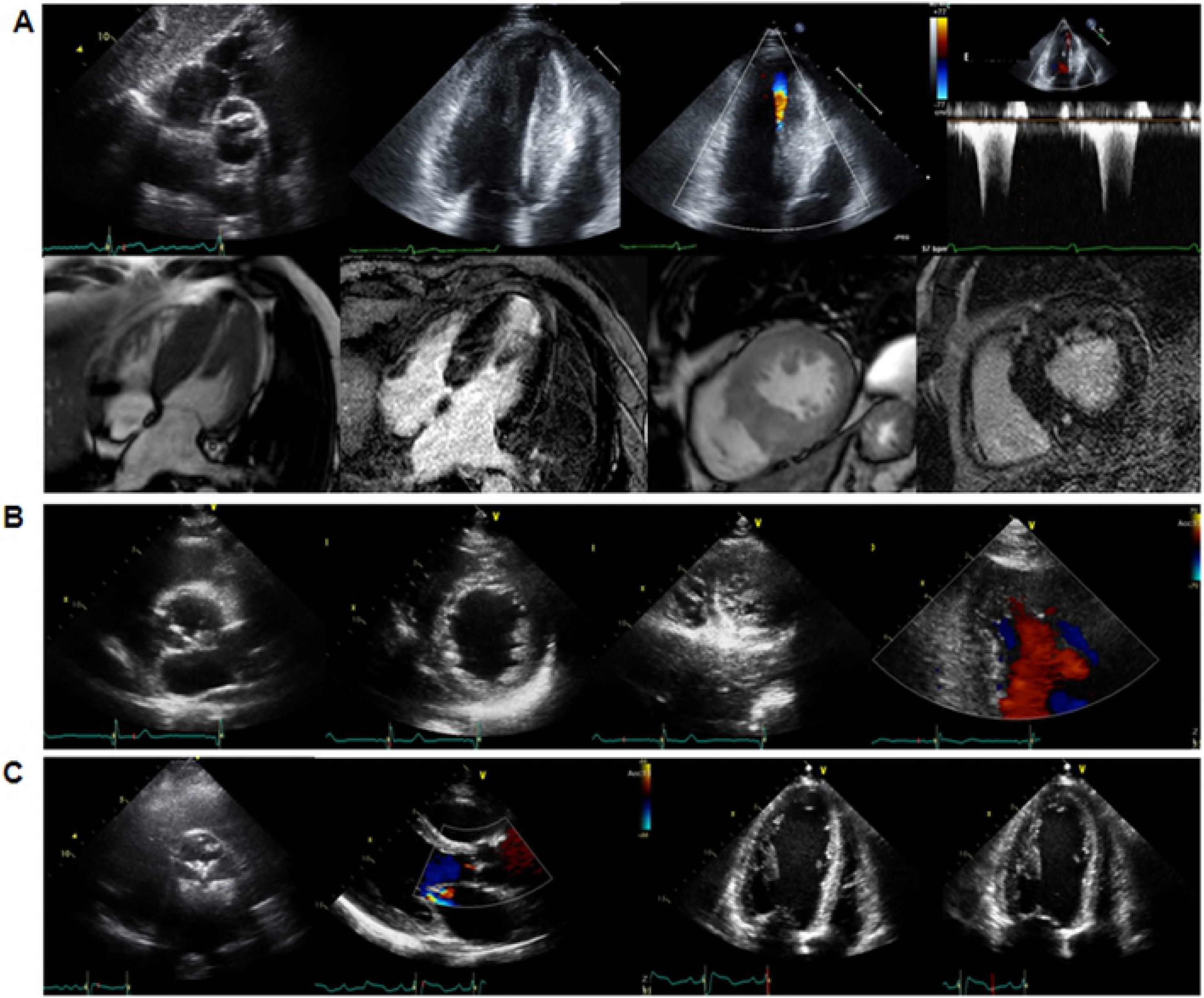
Representative cases of coexisting cardiomyopathies in BAV subjects (A) HCM (B) LVNC (C) DCM

### Clinical Significance of Concomitant CMs in BAV Subjects

Logistic regression analysis was performed to investigate the factors associated with heart failure in subjects with BAV. In univariate analysis, increased age was significantly associated with heart failure (odds ratio (OR) 1.035, p < 0.001), but gender was not. Comorbidities such as hypertension (OR 1.793, p < 0.001), diabetes mellitus (OR 1.704, p = 0.003), and chronic kidney disease (OR 2.226, p < 0.001) were significantly associated with heart failure. Aortic valve dysfunction was not a significant associating factor for heart failure. However, an increased aorta diameter (OR 1.020, p = 0.046) was significantly associated with heart failure. Moreover, presence of CM was the strongest factor associated with heart failure (OR 2.582, p < 0.001). In multivariate analysis, age, hypertension, and presence of concomitant CM were independently associated with heart failure (Table 4).

**Table 4.** Factors associated with heart failure in BAV subjects

| Variable | Univariate |  |  | Multivariate |  |  |
| --- | --- | --- | --- | --- | --- | --- |
|  | OR | 95% CI | P-value | OR | 95% CI | P-value |
| Age | 1.035 | 1.002-1.047 | <0.001 | 1.031 | 1.018-1.044 | <0.001 |
| Female | 1.164 | 0.835-1.623 | 0.370 |  |  |  |
| Body mass index | 1.027 | 0.986-1.069 | 0.197 |  |  |  |
| Hypertension | 1.793 | 1.320-2.435 | <0.001 | 1.547 | 1.124-2.128 | 0.007 |
| Diabetes mellitus | 1.704 | 1.196-2.429 | 0.003 |  |  |  |
| CKD | 2.226 | 1.420-3.489 | <0.001 |  |  |  |
| Significant AS or AR | 0.958 | 0.691-1.327 | 0.795 |  |  |  |
| Aortopathy | 1.320 | 0.973-1.790 | 0.074 |  |  |  |
| Aorta diameter | 1.020 | 1.000-1.040 | 0.046 |  |  |  |
| CM | 2.582 | 1.513-4.406 | 0.001 | 2.866 | 1.652-4.974 | <0.001 |

BAV, bicuspid aortic valve; OR, odds ratio; CI, confidence interval; CKD, chronic kidney disease; AS, aortic stenosis; AR, aortic regurgitation; CM, cardiomyopathy

## Discussion

The principal findings of the present study are as follows. First, approximately 6% of BAV subjects had concomitant CMs. LVNC was the most prevalent; however, that prevalence in BAV subjects was only 3.4% lower than the prevalence previously reported. Second, BAV subjects with specific CMs had distinct clinical and echocardiographic features. Third, age, hypertension, and presence of CM were significantly associated with heart failure. To the best of our knowledge, this is the first study in which the prevalence, characteristics, and clinical implications of concomitant specific CMs were identified in adults with BAV using a large Korean registry.

Case reports of specific CMs in BAV subjects have been sporadic; however, studies on their prevalence are limited. Agarwal et al.[12] reported the incidence of LVNC in patients with BAV. In their retrospective observational study, 12 of 109 BAV patients (11.0%) were diagnosed with LVNC based on echocardiographic criteria. The mean age at diagnosis was 33 ± 16.9 years. The incidence was greater and mean age at diagnosis was significantly lower than in our study. Chandra et al.[24] researched the incidence of LVNC in Caucasians and African-Americans but not Asians and revealed inter-racial difference in frequency in BAV patients. Therefore, the difference of incidence between the present study and Agarwal et al. [12]was likely due to the population characteristics including age and race. Padang et al. [13] reported the incidence of HCM in patients with BAV. In their retrospective cohort, 23 patients were diagnosed with HCM and BAV based on echocardiographic criteria. The mean age at diagnosis was 52 ± 16 years. Their study showed 0.9% of patients with HCM had coexistent BAV and 0.4% of patients with BAV had coexistent HCM. The incidence of BAV in the general population is usually approximately 1%; therefore, the incidence of BAV was not different between the HCM population and general population. In our study, 1.4% of patients had concomitant BAV and HCM, and these results are similar to a previous study. In the present study, 0.8% idiopathic DCM was identified in BAV subjects, similar to the results found in the general population. To date, studies on the prevalence of DCM in BAV subjects have not been published, probably because a certain degree of aortic valve dysfunction can result in chamber dilatation and myocardial dysfunction. To eliminate the confounding factors, patients with significant aortic valve dysfunction were excluded when diagnosing idiopathic DCM. Therefore, the actual prevalence of coexisting DCM in BAV subjects might be underestimated in this study. In a previous study, genetic factors were suggested as a possible cause of coexistent BAV and HCM[25]. However, few studies exist on genes in patients with BAV and specific CMs. In the future, genetic analysis will be important in studying the relationship between BAV and CM.

Recently, BAV has been recently noted not only in the valve itself, but also in relation to the left ventricle and aorta. Since BAV patients with concomitant CMs have prominent characteristics of their own myocardial disease, there were no significantly different characteristics in BAV morphology, aortic valve dysfunction, or aorta phenotype in overall group comparison. Although type 0 BAV phenotype and presence of aortopathy tended to be more prevalent in BAV patients with HCM, these tendencies did not show statistical significance.

Predictably, the presence of specific CMs may influence the patient’s clinical course, especially in aspects of heart failure in BAV subjects. Heart failure was associated with 16.9% of all BAV subjects and 32.8% of BAV subjects with specific CMs. In univariate analysis, age and well-known comorbidities were correlated with heart failure, as expected. Regarding several specific characteristics in BAV subjects, increased aortic diameter was positively associated with heart failure, but significant aortic valve dysfunction was not. These findings support a possible ventricular vascular interaction in BAV subjects, which has been suggested in previous studies [3, 7-9]. However, the strongest factor associated with heart failure was the presence of CM. Consequently, in multivariate analysis, age, hypertension, and presence of CM were independently associated with heart failure.

### Study Limitations

The present study had several limitations. First, this study included only Korean BAV subjects from a single tertiary referral center, which may result in bias. Therefore, multinational studies including various ethnic groups are needed to evaluate the prevalence of CMs in BAV subjects. However, we believe this study was the first report on the prevalence of concomitant CMs in a large Korean registry using comprehensive reviews. Second, data was lacking regarding common genetic backgrounds in BAV patients with CMs. The results of this study may be a basis for future genetic research. Third, aortic diameters were measured based on echocardiographic imaging alone because only some BAV subjects underwent computed tomography or cardiac magnetic resonance imaging.

## Conclusions

Concomitant BAV with CMs was observed in 5.6% of our BAV population. Several clinical and echocardiographic characteristics including comorbidities, heart failure presentation, BAV phenotypes, valve function, and presence of aortopathy were found in these patients. The presence of CM was independently associated with heart failure.

## Acknowledgement

None

## Supporting Information

### Author Contributions

Conceptualization: CYS.

Data curation: SYL HJJ.

Formal analysis: HJJ CYS.

Investigation: HJJ.

Methodology: CYS HJJ.

Project administration: CYS.

Resources: HJJ CYS DRK JYC KUC SYL GRH JWH.

Software: CYS HJJ.

Supervision: CYS.

Validation: CYS.

Visualization: HJJ CYS.

Writing - original draft: HJJ CYS.

Writing - review & editing: HJJ CYS.

## Abbreviations

BAV=: bicuspid aortic valve
LVNC=: left ventricular non-compaction
CM=: cardiomyopathy
HCM=: hypertrophic cardiomyopathy
DCM=: dilated cardiomyopathy
LVEF=: left ventricular ejection fraction
OR=: odds ratio
CI=: confidence interval

